# Effects of leaf removal on photosynthetic activity, fruit yield, and quality of micro-dwarf tomatoes

**DOI:** 10.64898/2026.05.10.724098

**Authors:** Dmitrii Usenko, Chen Giladi, Carmit Ziv, David Helman

## Abstract

Micro-dwarf tomato cultivars are increasingly considered for urban and controlled-environment agriculture due to their compact architecture and suitability for high-density planting. However, optimal canopy management strategies for these cultivars remain poorly defined. In this study, we evaluated the effects of different leaf removal intensities on leaf-level physiological performance, fruit yield, and fruit quality in three micro-dwarf tomato cultivars (*Mohammed, Hahms Gelbe Topftomate*, and *Red Robin*) grown under contrasting seasonal light conditions. Plants were subjected to low (15%), moderate (30%), or severe (90%) leaf removal, and leaf-level gas exchange was measured across canopy layers, along with yield and fruit quality assessments.

Severe leaf removal (90%) increased carbon assimilation, transpiration, and stomatal conductance in middle and lower canopy leaves by up to approximately twofold compared with control plants, indicating improved light availability at the leaf level. However, these physiological enhancements did not consistently translate into higher yield, reflecting reduced whole-plant source capacity under excessive leaf removal. Low to moderate leaf removal (15–30%) generally increased or maintained yield and fruit number, whereas severe leaf removal reduced yield in *Hahms Gelbe* and *Red Robin*, particularly under low seasonal radiation. In contrast, *Mohammed* exhibited yield increases of up to 220% under low leaf removal and maintained increased yield even under severe leaf removal under high-light conditions. Fruit quality was largely unaffected by leaf removal, except for total soluble solids, which declined by approximately 12% under severe leaf removal across cultivars, consistent with sugar dilution under source limitation.

Overall, these results demonstrate that optimal leaf removal in micro-dwarf tomatoes requires balancing improved canopy light distribution with maintenance of sufficient leaf area for carbon assimilation. Leaf removal thresholds are strongly cultivar- and light-dependent, emphasizing the need for cultivar-specific canopy management strategies in compact tomato systems and controlled-environment agriculture.

## 1. Introduction

Ongoing urbanization is increasing the demand for novel farming approaches tailored to food production within cities. These approaches include a range of technologies, such as multi-layered vertical farming systems (Harbinson & Taylor, 2025; van Delden et al., 2021). Certain vegetable cultivars are particularly well suited to such production environments. Among them, dwarf—and especially micro-dwarf—tomato cultivars are characterized by their compact plant architecture (Martí et al., 2006), resulting from a determinate growth habit that limits plant height.

Micro-dwarf tomatoes were initially developed as recreational plants; however, these same traits may make them highly suitable for urban agriculture. In addition, these cultivars exhibit relatively low light requirements (Kato et al., 2011), short production cycles (Sun et al., 2006), and tolerance to high planting densities, all of which are advantageous for cultivation in space-limited urban environments (Meissner et al., 1997). Despite these promising characteristics, micro-dwarf tomato cultivars remain far less studied for commercial production than standard indeterminate tomato types.

The work of Ke *et al*., (2022, 2023) shed light on the physiological characteristics of dwarf tomato cultivars. Specifically, they demonstrated that radiation-use efficiency (RUE) and fruit biomass radiation-use efficiency (FBRUE) increase with photosynthetic photon flux density (PPFD) up to the point at which the canopy’s light-absorptive capacity is saturated. Beyond this threshold, FBRUE declines as excess radiation exceeds the plant’s capacity to utilize additional energy.

In light of these findings, and given that dwarf tomato cultivars typically exhibit a bushy architecture with short internodes, partial leaf removal emerges as a potentially powerful tool for regulating canopy light interception and yield. Unlike standard indeterminate tomato cultivars, which require regular pruning to remove axillary shoots and limit excessive branching, compact determinate or semi-determinate cultivars generally do not require such interventions. Nevertheless, leaf removal is commonly practiced in these plants to improve light penetration (Appolloni et al., 2023), maintain adequate airflow within the canopy (Qin et al., 2011), and reduce disease pressure (Decognet et al., 2012)—an approach that may be particularly relevant in high-density, controlled-environment systems where within-canopy light distribution is a key constraint.

Empirical studies indicate that moderate defoliation can increase yield or, at a minimum, maintain it, whereas severe defoliation generally reduces yield. In tomato, removal of lower leaves has been shown to improve yield when the original canopy is overly dense; however, yield losses frequently occur when more than approximately 40% of the total leaf area is removed (Mondal, 2022; Saiful Islam et al., 2016; Slack, 1986; Stacey, 1983). Thus, the yield response to leaf removal depends on the intensity and timing of the treatment, the remaining total leaf area, and prevailing environmental conditions such as light availability and temperature. Notably, some studies have reported that removal of up to 60% of leaf area did not result in yield reduction, presumably due to compensatory increases in the photosynthetic rate of the remaining leaves (Raya et al., 2024).

From a physiological perspective, sugar production in tomato plants is generally sink-limited during early developmental stages and becomes source-limited during fruiting (Ho, 1996; Iqbal et al., 2012; Li et al., 2015). Leaf removal alters this balance by reducing total leaf area while potentially increasing photosynthetic productivity of leaves previously shaded. The threshold at which leaf removal becomes detrimental varies with light availability. For instance, under severe leaf removal, plants may be unable to produce sufficient assimilates to sustain desirable total soluble solids (TSS) concentrations in the fruit if the remaining leaf area is inadequate (Ho, 1996).

Within a source–sink framework, when carbon supply is constrained—as under excessive leaf removal—plants tend to prioritize overall fruit growth over sugar accumulation, resulting in dilution of soluble solids within the fruit flesh (Falchi et al., 2020). This trade-off arises because sink demand for biomass accumulation competes with solute accumulation, and reduced source strength limits the allocation of sugars to the fruit.

However, despite extensive research on leaf removal responses in standard tomato cultivars, the physiological, yield, and quality responses of micro-dwarf tomatoes—particularly under light conditions relevant to urban and controlled-environment agriculture—remain poorly understood.

In this study, we evaluate the effects of leaf removal on yield, carbon assimilation, transpiration, and fruit quality in three micro-dwarf tomato cultivars under two light condition regimes (seasons). The primary objective is to identify the advantages and limitations of different leaf removal intensities and to determine an optimal, balanced leaf removal strategy that maximizes yield while maintaining favorable plant physiological performance and high marketable fruit quality. Accordingly, this study addresses the following research questions:

1. *How does leaf removal intensity affect leaf-level physiological performance across different canopy layers in micro-dwarf tomato plants?*
2. *Does leaf removal enhance yield and fruit number in micro-dwarf tomato cultivars?*
3. *How does leaf removal intensity influence fruit quality traits in micro-dwarf tomatoes?*

## 2. Data and methods

### 2.1. Plant material

Three micro-dwarf tomato cultivars (Dwarf Tomato Project — A Co-Operative Venture) were selected for this study: *Mohammed, Hahms Gelbe Topftomate*, and *Red Robin* (Fig. 1).

**Figure 1.**
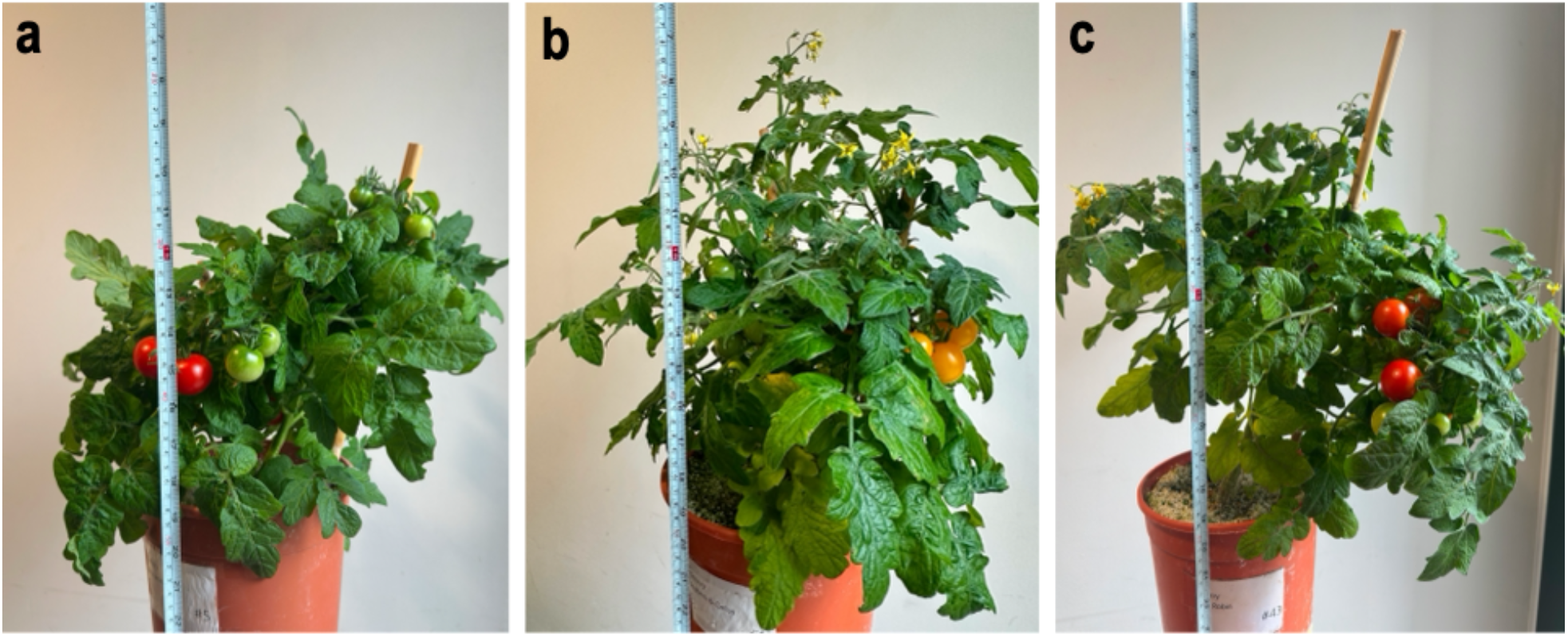
Pictures of **(a)** *Mohamed*, **(b)** *Hahms Gelbe Topftomate*, **(c)** *Red Robin* micro-dwarf tomato cultivars. The plants in the pictures have not had their leaves removed.

We chose these varieties for their compact growth habits and suitability for urban farming, particularly in hydroponic systems where space efficiency is crucial. All three dwarf tomato cultivars share a compact, determinate growth habit, making them well**-**suited for high-density planting and automated management systems. They produce cherry-sized tomatoes averaging 10– 12 grams each: *Mohammed* and *Red Robin* yield red fruits, while *Hahms Gelbe* produces bright yellow fruits, adding diversity to the dataset and potentially influencing light absorption. *Red Robin* is specifically known for its adaptability to confined spaces.

### 2.2. Glasshouse environmental conditions

Two experiments were conducted in a controlled glasshouse at the Robert H. Smith Faculty of Agriculture, Food and Environment, The Hebrew University of Jerusalem, Rehovot, Israel (31.90445°N/34.80459°E, Jiang *et al*., 2022; Mulero *et al*., 2023; Usenko *et al*., 2025) to assess the consistency and reliability of the method under varying seasonal light conditions. Other conditions—temperature, humidity, irrigation, and fertilizer application—were maintained similarly across both experiments, with daytime and nighttime temperatures and relative humidity held constant throughout (Table 1).

**Table 1.** Durations and growing conditions of the two experiments (*Exp. #1* and *Exp. #2*).

| Parameter | *Exp. #1* | *Exp. #2* |
| --- | --- | --- |
| *Timeline* | March 2023 — June 2023<br>(Spring to early summer) | September 2023 — December 2023<br>(Autumn to early winter) |
| *Temperature (fixed)* | 24 °C (daytime) / 18 °C (nighttime) |  |
| *Relative humidity* | 70–75% (daytime) / 90–97% (nighttime) |  |
| *Lighting conditions* | Natural sunlight. Average daily solar radiation was higher during this period, typical for Mediterranean spring and early summer (Fig. 2). | Natural sunlight. Shorter daylight hours and lower natural solar radiation due to seasonal changes (Fig. 2). |
| *Irrigation* | Plants were grown in 4 L pots filled with perlite, a soilless medium facilitating hydroponic growth. An automated drip irrigation system supplied 150 mL of water per plant five times daily, once in two hours, starting at 7:00 AM. |  |
| *Nutrition* | Nutrient solutions were provided in the irrigation water, containing balanced macro and micronutrients suitable for tomato cultivation. |  |
| *Pest control* | Integrated pest management strategies, including regular inspections and pesticide application, were employed to prevent infestations that could affect plant growth and leaf area. |  |

Since the glasshouse lacks artificial lighting, changes in natural solar radiation throughout the season are likely to affect plant growth and yields. Spring and summer in this area benefit from increased natural sunlight, promoting robust vegetation growth and potentially larger leaf areas, whereas autumn and winter receive less sunlight. Accordingly, the first experiment (*Exp. #1*) was run during the spring and summer and radiation levels consistently peaked between 800 and 1,000 μmol m^−2^ s^−1^, while the second experiment (*Exp. #2*) was conducted in autumn and winter, when radiation levels decrease (Fig. 2). While the experiment began in September with light levels comparable to *Exp. #1* (around 850 μmol m^−2^ s^−1^), in late November and December, the shorter day lengths and the sun’s lower position in the sky caused PAR levels to drop to an average weekly maximum of under 400 μmol m^−2^ s^−1^ (Fig. 2).

**Figure 2.**
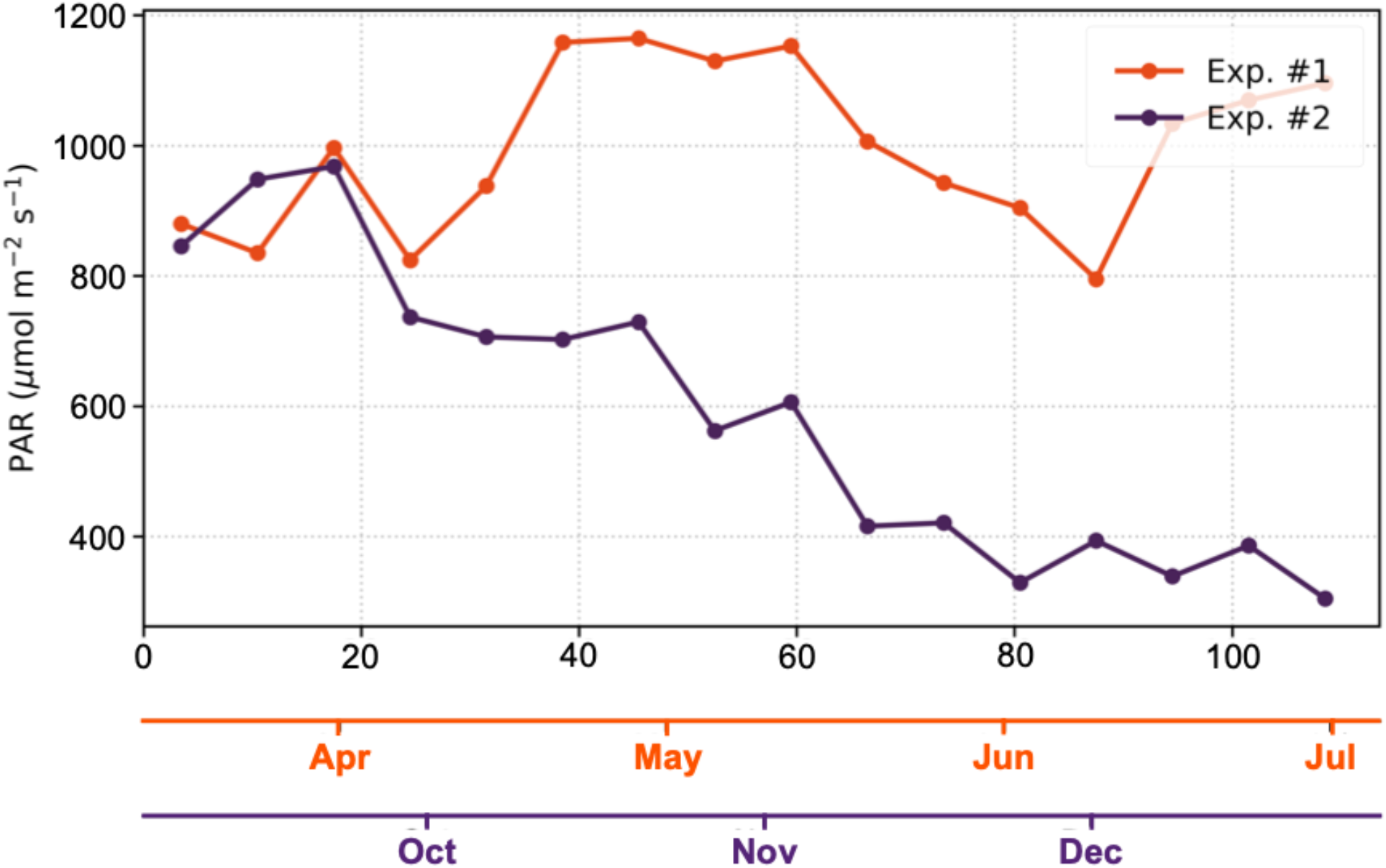
Average weekly maximum photosynthetically active radiation (PAR) levels during the 110 days of the experiments and their corresponding growing seasons (*Exp. #1* from March to June 2023 in orange and *Exp. #2* from September to December 2023 in dark purple).

### 2.3. Experimental design

A set of 60 plants from three cultivars (20 per cultivar) was used in each experiment. Plants were grown from seeds purchased at the LittleIslandSeedCo shop on Etsy (Canada). The seeds were planted on Feb 1, 2023, in *Exp. #1* and on July 24, 2023, in *Exp. #2* in disposable planters filled with 80/20 soil and perlite mixture. The planters were topped with thin layers of vermiculite and covered with paper pending germination. All plants were transplanted in 4 mm perlite on March 15, 2023, in *Exp. #1*, and on September 6, 2023, in *Exp. #2* in 4L pots.

The plants were then evenly divided into four leaf-removal treatment groups (Fig. 3). A control group (C), with plants undergoing no deliberate defoliation, except the routine removal of dry or damaged leaves; low-leaf removal (LR) treatment, with approximately 15% of the total leaves removed from the lower part of the plants (LR15%); a moderate LR treatment, with approximately 30% of the basal leaves removed (LR30%); and a severe-LR treatment, with nearly 90% of the leaves removed (LR90%) as follows, all leaves located within the bottom 30% of the plant height were removed, along with all upper-canopy leaves that were not directly adjacent to flowers or fruits. The LR90% treatment was intentionally designed as an extreme leaf removal scenario to identify physiological and productivity thresholds under severe source limitation.

**Figure 3.**
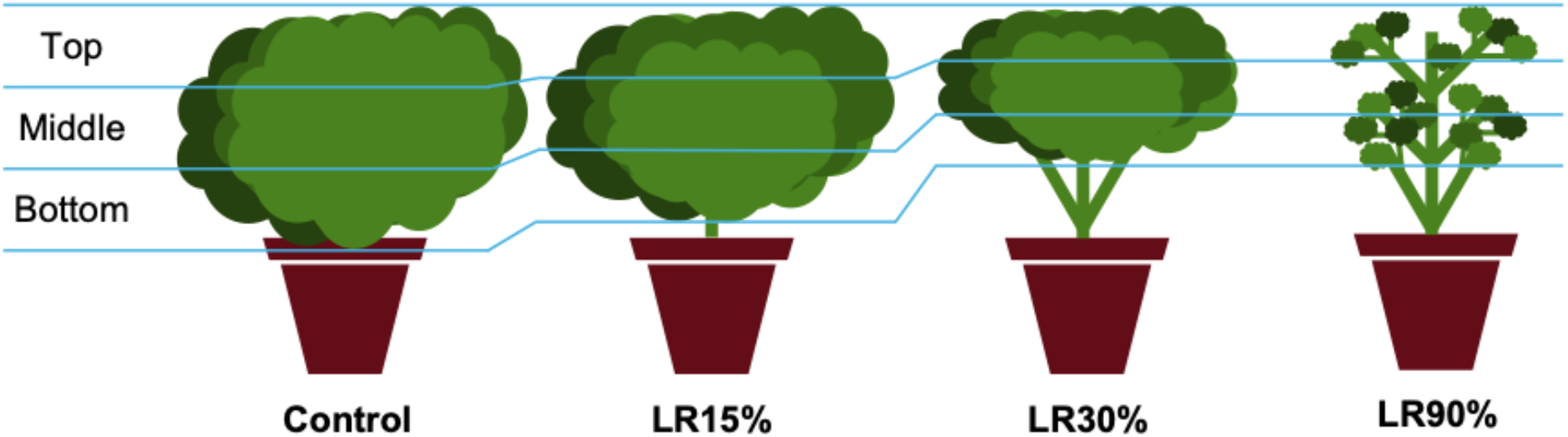
The leaf removal treatments, showing no leaf removal (Control), and leaf removal of various intensities, with low (LR15%), moderate (LR30%), and severe (LR90%) leaf removals from the bottom of the plants (except for LR90%, which also included leaf removal from the upper canopy). The scheme for analyzing photosynthetic activity across the top, middle, and bottom canopy layers is also shown.

Leaf removal was conducted in three discrete stages corresponding to key phenological phases: at the onset of flowering, at the beginning of fruit formation, and at peak plant productivity, occurring on days 22, 44, and 83 of the experiments, respectively.

The plants were distributed in the glasshouse according to a randomized complete block design to eliminate the effect of the radiation gradient.

### 2.4. Measurements

During the experiments, specific measurements were taken to estimate plants’ photosynthetic activity, productivity, and fruit quality. The measurements were taken at similar timings across the two experiments when possible.

Figure 4 outlines the different measurements and their timings per experiment.

**Figure 4.**
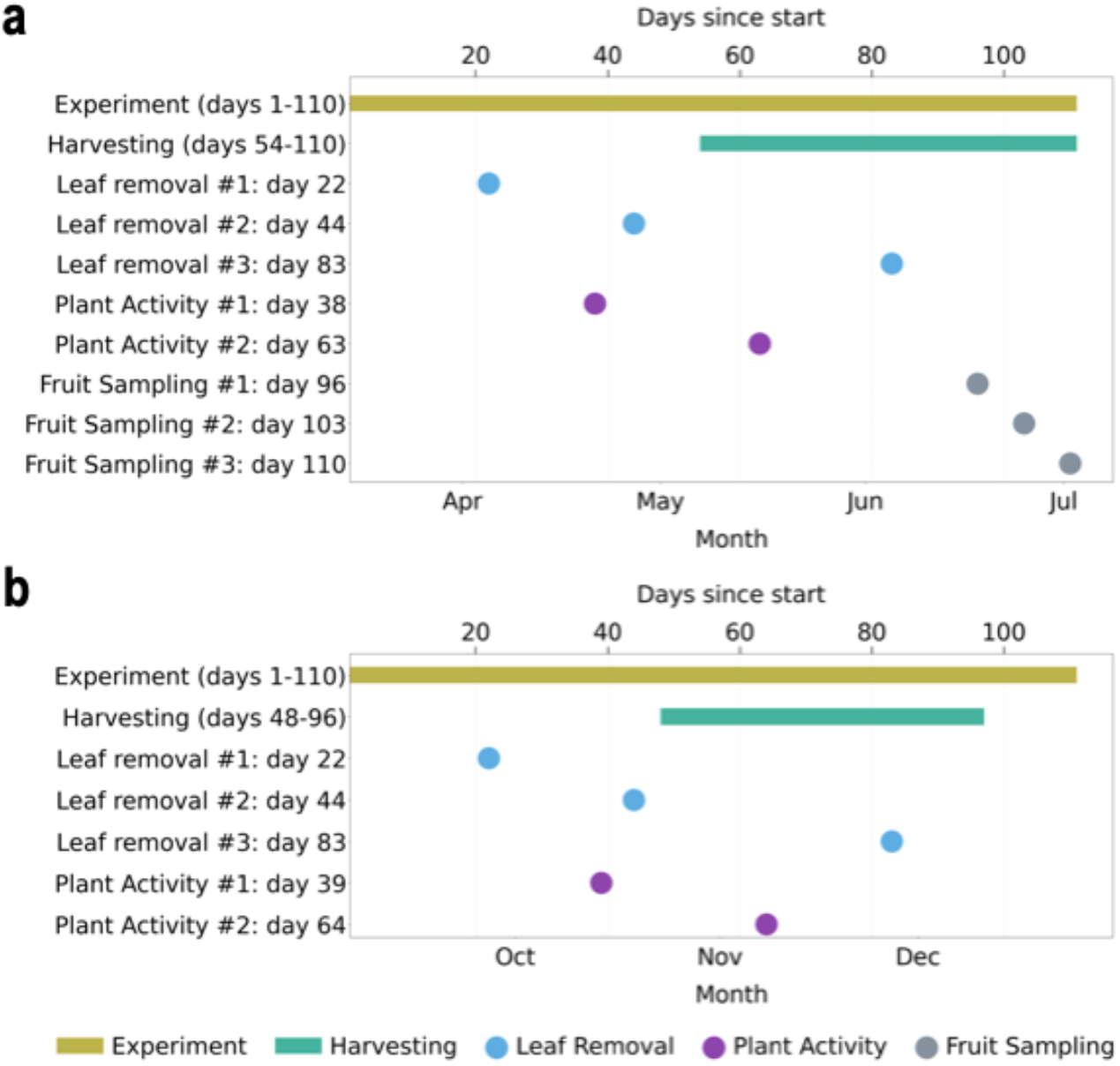
Timelines of the two experiments and measurements: **(a)** *Exp. #1* was conducted in the spring and summer of 2023, and **(2)** *Exp. #2* was conducted during the autumn and winter of 2023. Fruit quality sampling was conducted only in *Exp. #1*.

#### 2.4.1. Plant physiological activity

Plant photosynthetic activity was assessed using an infrared gas analyzer (LI-6800, Licor, Lincoln, USA). The gas analyzer was used to measure the carbon assimilation (*A*_n_; μmol m^-2^ s^-1^), transpiration (*T*_r_; mmol m^-2^ s^-1^), and stomatal conductance (*g*_s_; mol m^-2^ s^-1^). Selected plants were measured twice in each experiment: first at the beginning of the fruit-formation stage and second during peak productivity (days 38 and 63 in *Exp. #1*, and days 39 and 64 in *Exp. #2*). The measurements were conducted between 10 am and 12 pm under sunny conditions. Three leaves were chosen from the selected plants: one from the bottom, one from the middle, and one from the top (Fig. 3). All measured leaves were fully expanded and exposed to sunlight.

The fan speed was set to 10,000 rpm and the flow rate to 500 µmol s^−1^ in the measurement chamber. RH and temperature were left uncontrolled and therefore reflected ambient glasshouse conditions. Photosynthetic photon flux density (PPFD) was maintained at 1200 µmol m^−2^ s^−1^, and the reference CO_2_ level was set to 400 ppm (ambient conditions). The practice of recording measurements after stabilizing the gas exchange parameters was followed, which took around 1-2 minutes (Jiang et al., 2022).

#### 2.4.2. Yield

Fruits were collected from the plants when fully ripe (manually assessed by color and firmness). Each fruit was then weighed, and the harvest date and the total number of fruits per harvest were recorded.

#### 2.4.3. Fruit quality

Chemical analyses were performed to determine the following fruit quality parameters: total soluble solids (TSS), citric acid content, ascorbic acid content, and pH.

Fruits were sampled from the three cultivars in control groups and under treatments across three consecutive sampling dates: 18 June, 25 June, and 2 July 2023 (96, 103, and 110 days from the beginning of *Exp. #1*). For all cultivars, six control plants were sampled on each of the three dates. For *Hahms Gelbe*, the treatment groups each comprised six plants, collected on the first sampling date (18 June; 96 days from the beginning of the experiment). For *Red Robin*, each treatment group comprised six plants collected on the second sampling date (25 June; 103 days from the beginning of the experiment). For *Mohammed*, each treatment group comprised six plants collected on the final sampling date (2 July; 110 days from the beginning of the experiment).

These samples included both ripe and green fruits (Fass et al., 2025), but in the current study, only results for ripe fruits were analyzed and presented.

Fruits were stored at –80°C after collection. On the day of the tests, the fruits were unfrozen, and their flesh was filtered through 3 layers of sterile gauze. This juice was used for the tests.

For TSS (°Brix), we used a digital refractometer (Atago, Japan). The juice was pipetted into the chamber and measured three times. The average value was derived.

For citric acid (%) and pH, we used an 855 Robotic Titrosampler (Ω Metrohm, Dr. Golik Scientific Solutions, Israel). To prepare samples for this test, 1 mL of juice was diluted in 40 mL of double-distilled water. The titration device was calibrated using 1% citric acid standard solution.

Ascorbic acid (mg/100g) was measured at room temperature using the Folin reagent method (Jagota & Dani, 1982). 200 mg of fruit tissue was mixed with 0.8 mL of 10% trichloroacetic acid (Kim et al., 1987) and centrifuged at 13,000 rpm for 10 minutes. Then, 160 μL of DDW, 40 μL of supernatant, and 20 μL of 10-fold-diluted Folin reagent were mixed in a 96-well plate. Absorbance was measured at 760 nm after a 10-minute incubation in the dark. The ascorbate concentration was finally calculated from an L-ascorbic acid standard curve.

### 2.5. Statistical analyses

Data was analyzed using 3-way and 4-way Analysis of Variance (ANOVA), followed by Tukey’s Post-Hoc HSD and Dunnett’s tests, only after confirming normal data distribution through a Shapiro—Wilk test. HC3-robust version of Dunnett’s test was used when appropriate, while p-values were Holm-adjusted. All tests were run with α=0.05 (two-sided). The tests were run in JMP 15 Pro (© SAS Institute, Inc.) and via custom Python scripts using statsmodels (Seabold & Perktold, 2010) and SciPy (Virtanen et al., 2020) packages.

## 3. Results

### 3.1. Leaf-level photosynthesis, transpiration, and stomatal conductance

Cultivars did not differ in leaf carbon assimilation (*A*_n_), transpiration (*T*_r_), or stomatal conductance (*g*_s_) rates across experiments (no significant ‘Cultivar’ effect or ‘Cultivar × Experiment’ interaction in Table 2). *A*_n_, *T*_r_, and *g*_s_ were all sensitive to leaf removal. Yet, this effect was significant only for LR90% and only in leaves at the middle and bottom layers (‘Leaf position’ in Tables 2 and 3). LR90% increased gas-exchange rates in middle and bottom leaves in both experiments, with statistically significant increases for *g*_*s*_, *A*_n_ in middle leaves in *Exp. #1*, and *T*_r_ in the bottom leaves (in both experiments), and the middle leaves only in *Exp. #1* (Tables 2 and 3).

**Table 2.**
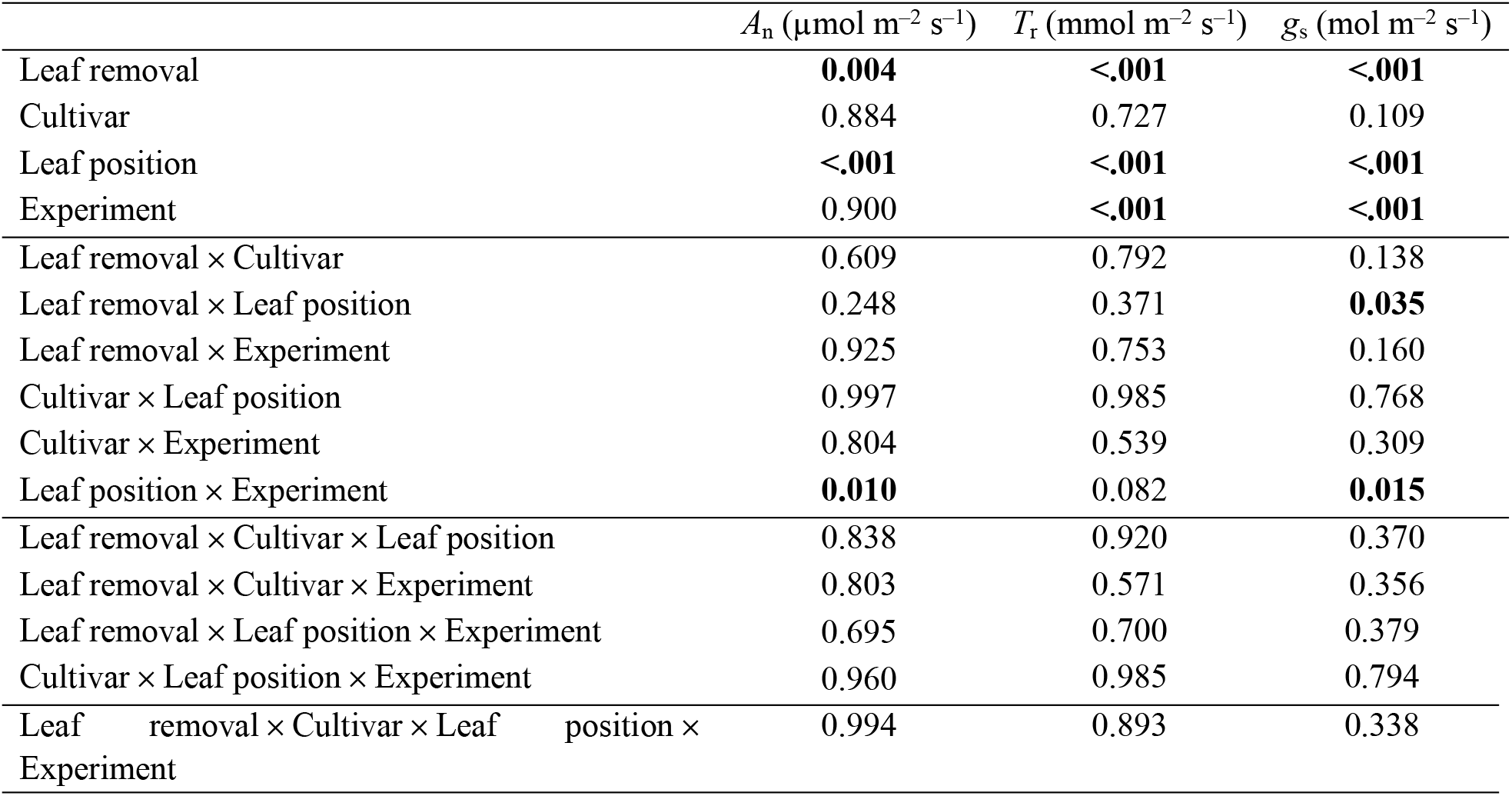
Effect estimations (*p*-values) for leaf-level carbon assimilation (*A*_n_), transpiration (*T*_r_), and stomatal conductance (*g*_s_) based on a 4-way ANOVA. Highlighted in bold are statistically significant effects (*p*<0.05).

**Table 3.** Mean values (±1σ) of leaf-level carbon assimilation (*A*_n_), transpiration (*T*_r_), and stomatal conductance (*g*_s_) rates per treatment and leaf position for *Exp. #1* and *Exp. #2*. Highlighted in bold are the treatments that differ significantly from the control group according to Dunnett’s test, calculated separately for each leaf position across the two experiments.

| Leaf position | Treatment | $A_n$ ($\mu\text{mol m}^{-2} \text{s}^{-1}$) | | $T_r$ ($\text{mmol m}^{-2} \text{s}^{-1}$) | | $g_s$ ($\text{mol m}^{-2} \text{s}^{-1}$) | |
| --- | --- | --- | --- | --- | --- | --- | --- |
|  |  | Exp. #1 | Exp. #2 | Exp. #1 | Exp. #2 | Exp. #1 | Exp. #2 |
| Top | C | 18.6 ± 4.2 | 15.8 ± 8.7 | 7.2 ± 1.4 | 12.1 ± 4.3 | 0.62 ± 0.15 | 0.79 ± 0.29 |
|  | LR15% | 17.9 ± 4.8 | 16.3 ± 7.6 | 7.2 ± 1.5 | 12.2 ± 4.5 | 0.58 ± 0.17 | 0.76 ± 0.13 |
|  | LR30% | 16.9 ± 4.0 | 14.2 ± 7.7 | 7.9 ± 1.1 | 11.2 ± 5.2 | 0.61 ± 0.13 | 0.71 ± 0.26 |
|  | LR90% | 19.7 ± 5.5 | 16.0 ± 8.3 | 8.3 ± 1.3 | 12.1 ± 4.6 | 0.66 ± 0.15 | 0.85 ± 0.22 |
| Middle | C | 6.4 ± 5.1 | 9.9 ± 8.2 | 3.0 ± 2.5 | 10.1 ± 5.0 | 0.21 ± 0.21 | 0.55 ± 0.20 |
|  | LR15% | 7.7 ± 5.4 | 10.1 ± 5.9 | 3.5 ± 2.3 | 10.2 ± 4.7 | 0.25 ± 0.21 | 0.54 ± 0.15 |
|  | LR30% | 4.6 ± 4.2 | 9.7 ± 5.4 | 2.5 ± 2.0 | 8.8 ± 5.5 | 0.14 ± 0.13 | 0.47 ± 0.26 |
|  | LR90% | **13.5 ± 3.0** | 13.5 ± 7.6 | **6.2 ± 1.6** | 11.2 ± 4.0 | **0.43 ± 0.17** | **0.73 ± 0.21** |
| Bottom | C | 4.8 ± 4.4 | 4.9 ± 7.3 | 2.2 ± 2.0 | 6.2 ± 3.3 | 0.13 ± 0.15 | 0.29 ± 0.17 |
|  | LR15% | 5.0 ± 3.8 | 4.6 ± 5.7 | 2.5 ± 1.9 | 6.5 ± 4.7 | 0.15 ± 0.15 | 0.29 ± 0.19 |
|  | LR30% | 7.3 ± 3.5 | 5.8 ± 6.9 | 3.0 ± 1.6 | 5.9 ± 3.5 | 0.16 ± 0.10 | 0.27 ± 0.17 |
|  | LR90% | 7.8 ± 3.7 | 10.4 ± 7.4 | **4.6 ± 2.2** | **10.6 ± 4.1** | **0.29 ± 0.16** | **0.66 ± 0.26** |

### 3.2. Fruit yield and number

Plants were more productive in *Exp. #1* than in *Exp. #2* (*p*<0.001 from a two-sided *t*-test; Figs. 5a and 6a), as expected from higher radiation during the spring-summer period. Their harvesting period started earlier (Figs. 5b,c and 6b,c), and their production rate was higher than that in *Exp. #2* throughout the season (a steeper increase in Figs. 5b,c, and 6b,c).

**Figure 5.**
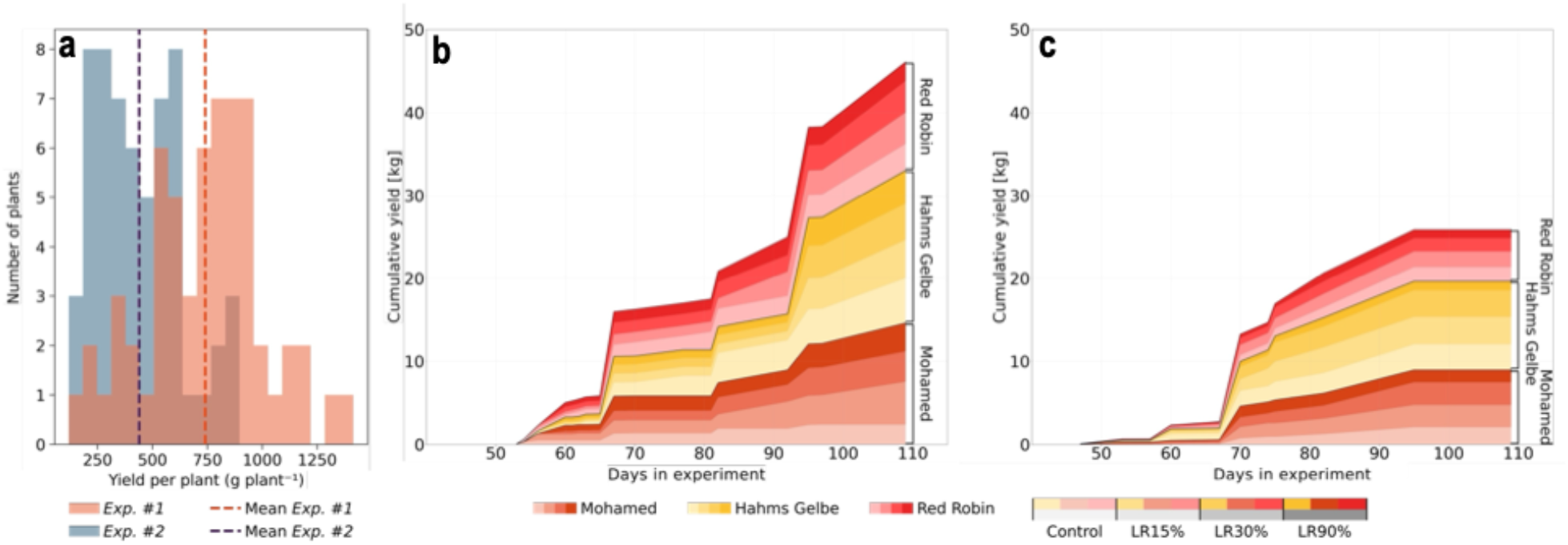
Yield distributions of the two experiments (*Exp. #1* and *Exp. #2*). (**a**) The overall distribution for all harvest periods, and their distribution over time in **(b)** *Exp. #1* and **(c)** *Exp. #2*. Areas of different hues indicate cultivars within treatments, with the least saturating color representing no leaf removal, and the most saturated color representing 90% leaf removal.

**Figure 6.**
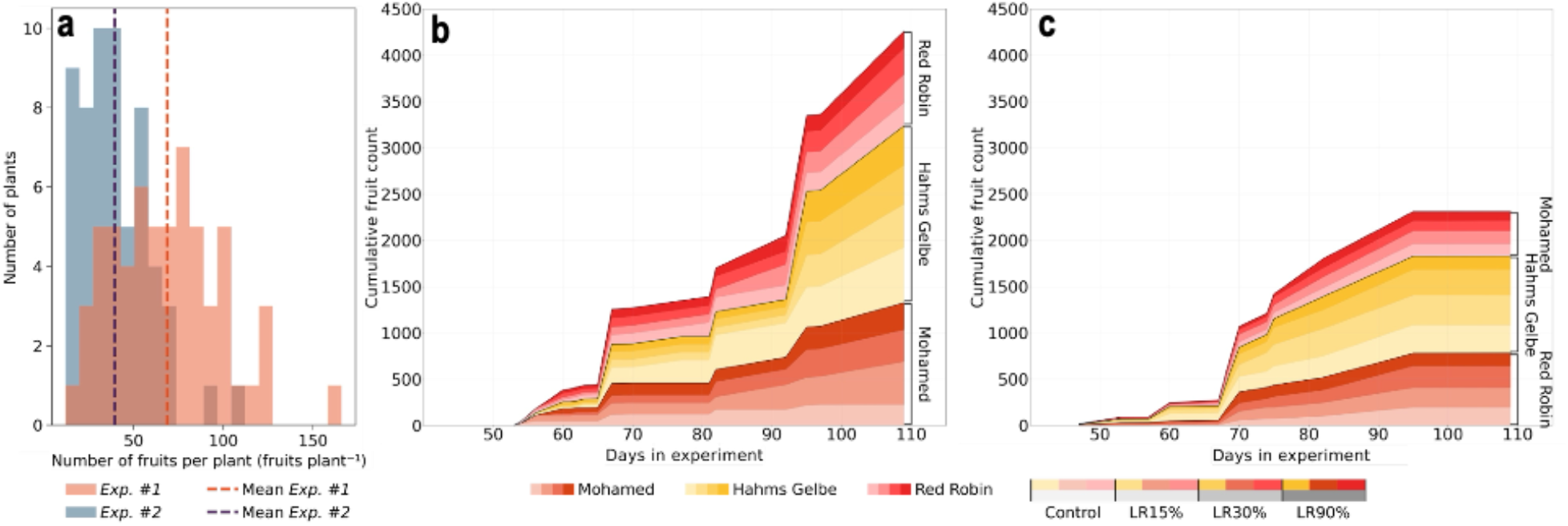
Fruit quantity distributions of the two experiments (*Exp. #1* and *Exp. #2*). (**a**) The overall distribution for all harvest periods, and their distribution over time in **(b)** *Exp. #1* and **(c)** *Exp. #2*. Areas of different hues indicate cultivars within treatments, with the least saturating color representing no leaf removal, and the most saturated color representing 90% leaf removal.

Yield and number of fruits were both highest in *Hahms Gelbe* (910.9±184.1 g plant^−1^ and 94.7±23.8 fruits plant^−1^ in *Exp. #1*; and 539.6±224.2 g plant^−1^ and 53.5±20.7 fruits plant^−1^ in *Exp. #2*) and lowest in *Red Robin* (619.3±232.4 g plant^−1^ and 49±19.7 fruits plant^−1^ in *Exp. #1*; and 310.8±99.1 g plant^−1^ and 24.2±7.9 fruits plant^−1^ in *Exp. #2*) and *Mohammed* (693.1±310.2 g plant^−1^ and 63.1±27 fruits plant^−1^ in *Exp. #1*; and 471.6±166 g plant^−1^ and 41±14.3 fruits plant^−1^ in *Exp. #2*).

Leaf removal affected both fruit yield and number, with the effect on yield depending on cultivar and seasonal light conditions (‘Leaf removal × Cultivar’ and ‘Leaf removal × Experiment’ significant interactions in Table 4). The effect of light conditions on the number of fruits was only marginally significant (*p*=0.075 for ‘Leaf removal × Experiment’).

**Table 4.** Effect estimations (*p*-values) for fruit yield and number from a 3-way ANOVA. Highlighted in bold are the significant effects (*p*<0.05).

| Factor | Yield (g plant<sup>-1</sup>) | Fruit number (# plant<sup>-1</sup>) |
| --- | --- | --- |
| Leaf removal | **<.0001** | **<.0001** |
| Cultivar | **<.0001** | **<.0001** |
| Experiment | **<.0001** | **<.0001** |
| Leaf removal × Cultivar | **<.0001** | **<.0001** |
| Leaf removal × Experiment | **0.005** | 0.075 |
| Cultivar × Experiment | 0.066 | **0.013** |
| Leaf removal × Cultivar × Experiment | **0.001** | **0.003** |

In general, LR90% had a significant adverse effect on the yield of all three cultivars, with a statistically significant negative effect only on *Hahms Gelbe* and *Red Robin*, and only under low light conditions (*Exp. #2*; Fig. 7b). The exception was *Mohammed*, which showed increased yield even under intense leaf removal, but only under high light conditions (*Exp. #1*; Fig. 7a). The most substantial increase in *Mohammed*’s yields, however, occurred under low and moderate leaf removal treatments, with +708.9 and +414.2 g plant^−1^, corresponding to yield increases of ∼220% and ∼128% for LR15% and LR30%, respectively. A similar leaf removal effect, yet less pronounced than that of yield, was observed on fruit number (Fig. 7c,d).

**Figure 7.**
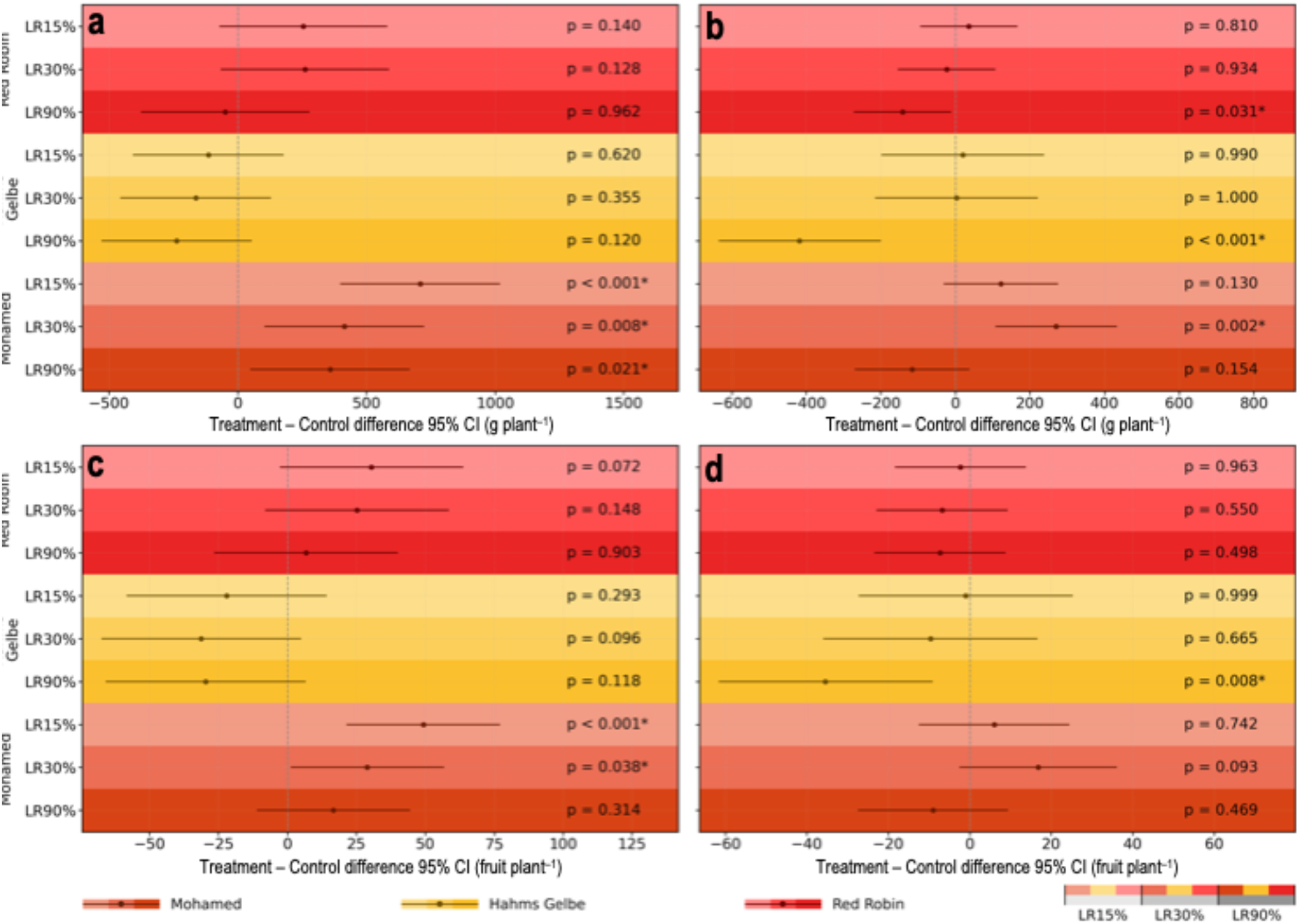
Results of Dunnett’s test for leaf removal treatments effect on (**a, b**) yield and (**c, d**) fruit number per cultivar in *Exp. #1* (**a, c**) and (**b, d**) *Exp. #2*.

### 3.3. Fruit quality

Cultivars did not differ in fruit quality (no ‘Cultivar’ effect in Table 5), while leaf removal affected TSS, regardless of the cultivar (no significant ‘Leaf removal × Cultivar’ interaction in Table 5). The pH, total soluble solids (TSS), and ascorbic acid contents of the fruits were generally within the ranges typically reported for cherry tomatoes, whereas citric acid content was slightly higher, varying between 0.70 and 0.77% (Table 6).

**Table 5.**
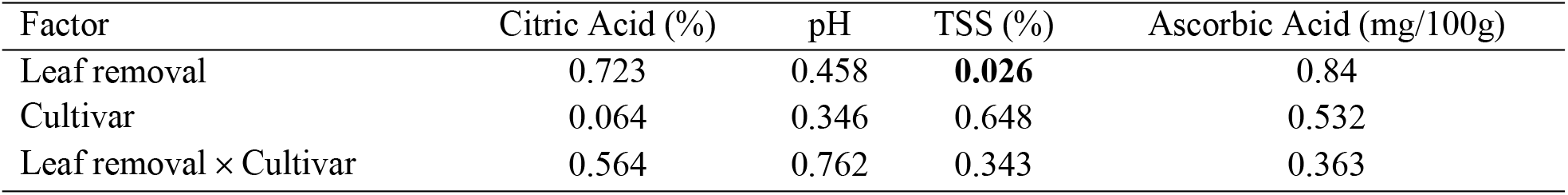
Effect estimations for fruit quality (citric acid, pH, total soluble solids, and ascorbic acid) based on a two-way ANOVA. Significant effects (*p*<0.05) are highlighted in bold.

**Table 6.**
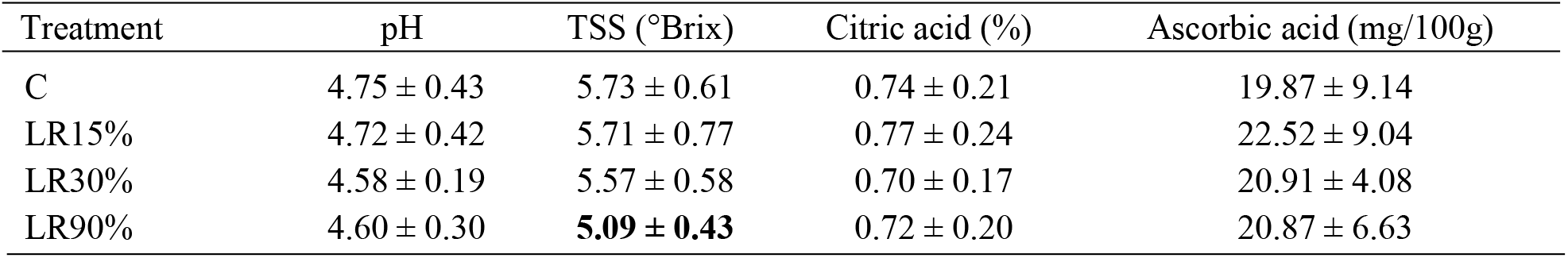
Mean values (±1σ) of pH, total soluble solids (TSS), citric acid content, and ascorbic acid content. Highlighted in bold are the treatments that differ significantly from the control group according to Dunnett’s test.

Severe leaf removal (LR90%) reduced TSS in fruits by ∼12%, while more moderate LRs (LR15% and LR30%) did not significantly affect TSS (Fig. 8). pH, citric, and ascorbic acids were unaffected by leaf removal, while seasonal effects were not assessed because quality parameters were collected only in *Exp. #1* (spring-summer light conditions).

**Figure 8.**
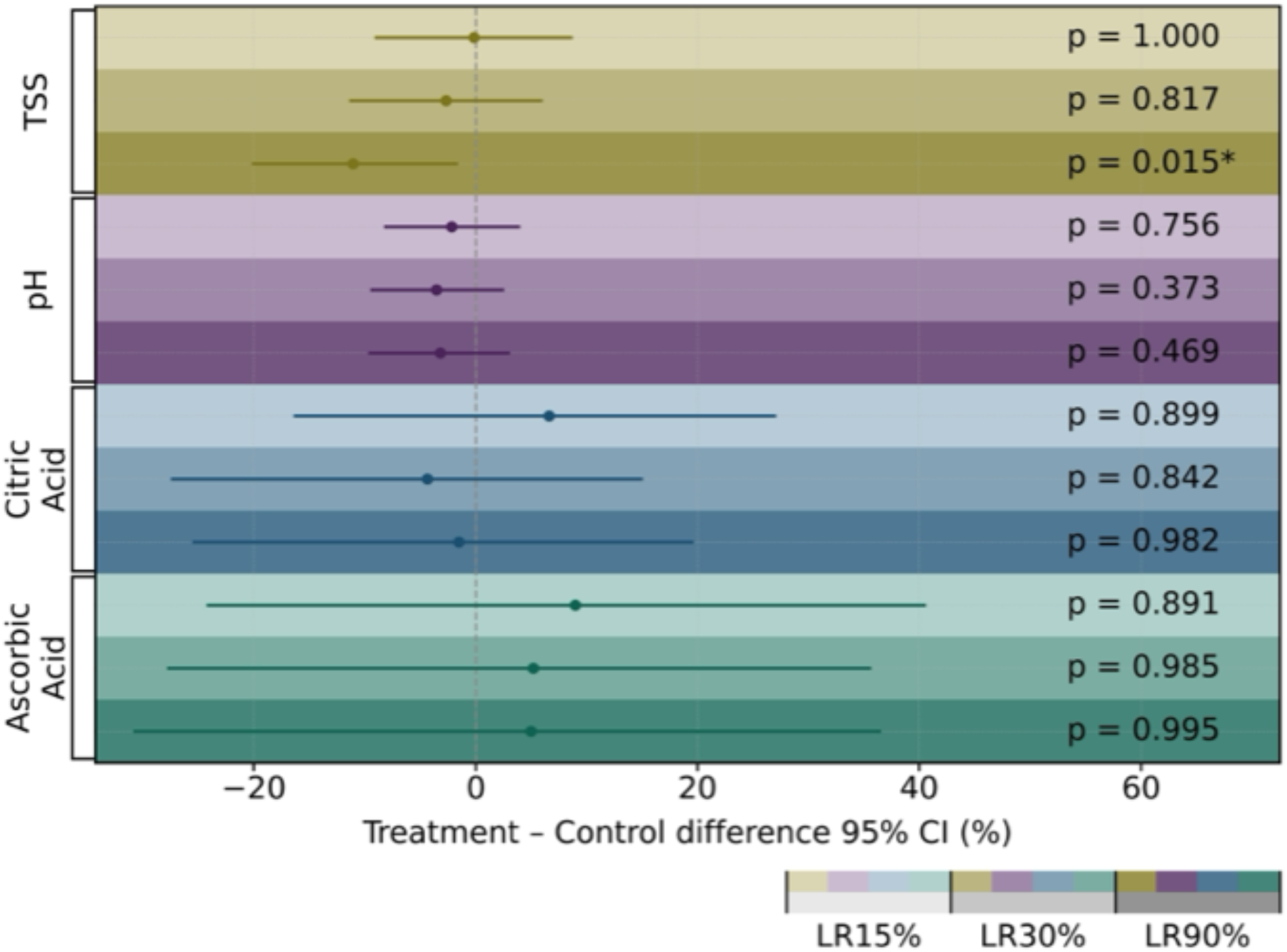
Leaf removal effects on fruit quality (TSS, pH, citric acid, and ascorbic acid). The *p*-values represent the results of a Dunnett’s test.

## 4. Discussion

The results of this study demonstrate that the effects of leaf removal on micro-dwarf tomato performance are strongly dependent on removal intensity, cultivar, and seasonal light conditions. While leaf removal altered leaf-level physiological performance in predictable ways, its effects on yield and fruit quality reveal important constraints imposed by compact canopy architecture and source–sink relationships in micro-dwarf tomatoes.

### 4.1. Leaf removal alters leaf-level physiology but does not necessarily enhance whole-plant carbon gain

Leaf removal affected leaf-level gas exchange primarily in the middle and lower canopy layers, and only under severe leaf removal (LR90%). In these treatments, carbon assimilation, transpiration, and stomatal conductance increased in remaining leaves, whereas top-canopy leaves maintained relatively stable gas-exchange rates across treatments. This pattern suggests that severe leaf removal alleviated self-shading and improved light availability for previously light-limited leaves, thereby enhancing their photosynthetic activity.

However, increased photosynthetic efficiency at the individual-leaf level did not translate into improved whole-plant productivity in most cultivars. This highlights a key physiological trade-off in compact canopies: while severe leaf removal can improve light penetration and leaf-level photosynthesis, it simultaneously reduces total leaf area and thus the canopy’s overall capacity for radiation interception and carbon assimilation. Consequently, enhanced per-leaf performance may be offset—or even outweighed—by reductions in total source strength.

This decoupling between leaf-level physiology and whole-plant performance is consistent with previous findings in dwarf tomato systems, where radiation-use efficiency declines once light availability exceeds the canopy’s absorptive or biochemical capacity (Ke et al., 2022, 2023). Our results emphasize that, in compact micro-dwarf canopies, maintaining sufficient leaf area is critical for sustaining whole-plant carbon gain, even when individual leaves operate more efficiently.

### 4.2. Yield responses reveal cultivar-specific optima and source limitations under severe leaf removal

Low and moderate leaf removal (LR15% and LR30%) generally increased or maintained yield and fruit number in two of the three cultivars, confirming that limited leaf removal can be beneficial in compact tomato canopies. These treatments likely improved light distribution within the canopy without substantially compromising total leaf area, thereby optimizing the balance between radiation interception and photosynthetic efficiency.

In contrast, severe leaf removal (LR90%) caused pronounced yield reductions in most cultivars, particularly under low-light conditions (Exp. #2). This aligns with previous studies showing that excessive defoliation reduces yield once the reduction in source capacity outweighs gains in light penetration and gas exchange (Mondal, 2022; Raya et al., 2024; Saiful Islam et al., 2016). Under low seasonal radiation, this threshold appears to be reached more rapidly, rendering plants especially vulnerable to source limitation.

Notably, the cultivar *Mohammed* deviated from this general pattern, exhibiting increased yield even under severe leaf removal during the high-light spring–summer season. This suggests a cultivar-specific response that may reflect differences in canopy architecture, source–sink balance, or intrinsic fruit sink strength. One possible explanation is that *Mohammed* may be relatively sink-limited under control conditions, such that severe leaf removal reduces vegetative competition and reallocates assimilates toward fruit growth when light availability is not limiting. Alternatively, differences in leaf orientation or canopy density may allow *Mohammed* to benefit more from improved light penetration following leaf removal. While these hypotheses cannot be fully resolved within the scope of this study, they highlight the importance of genetic background in determining optimal leaf removal strategies for micro-dwarf tomatoes.

### 4.3. Fruit quality responses indicate sugar dilution under severe source limitation

Fruit quality was largely unaffected by leaf removal, with the exception of total soluble solids (TSS), which declined significantly only under severe leaf removal. The absence of changes in fruit size (*p*>0.1 for all LR treatments from Dunnett’s test; not shown) alongside reduced TSS from 5.73 ± 0.61 °Brix in control plants to 5.09 ± 0.43 °Brix in LR90% suggests that this effect reflects sugar dilution rather than impaired fruit growth.

From a source–sink perspective, severe leaf removal likely constrained assimilate supply during fruit development, while sink demand for biomass accumulation remained relatively stable. Under such conditions, plants tend to prioritize fruit growth over solute accumulation, leading to reduced sugar concentration in the fruit flesh (Falchi et al., 2020; Ho, 1996). The sensitivity of TSS to severe leaf removal, even when yield effects were cultivar- and season-dependent, underscores that fruit quality may be a more sensitive indicator of source limitation than yield alone in compact tomato systems.

### 4.4. Implications for micro-dwarf tomato management in controlled environments

Together, these findings indicate that management strategies developed for standard indeterminate tomato cultivars cannot be directly transferred to micro-dwarf tomatoes. In compact architectures, excessive leaf removal rapidly induces source limitation, even when leaf-level photosynthetic rates increase. Low-to-moderate leaf removal can improve canopy structure and productivity, but optimal removal thresholds are both cultivar-specific and light-dependent. Different canopy management approaches, including intra-canopy illumination, can show similar, but more consistent effects in bell pepper (Tiwari et al., 2022; Kamara et al., 2023).

These insights are particularly relevant for urban and controlled-environment agriculture, where high planting densities and fixed light regimes exacerbate canopy self-shading. In such systems, carefully calibrated leaf removal can be a valuable tool for optimizing light distribution and radiation-use efficiency, provided that sufficient leaf area is maintained to support whole-plant carbon assimilation. Future work should examine whether cultivar-specific architectural traits can be leveraged to refine leaf-removal strategies and improve productivity in space-limited growing systems. These findings support the development of cultivar-specific canopy management strategies for micro-dwarf tomatoes grown in controlled environments.

## Acknowledgments

The authors thank Yedidya Harris for maintaining the greenhouse plants and Gabriel Mulero for assisting with data collection and measurements. D.U. acknowledges personal funding from the SCE-HUJI Scholarship Program for Ph.D. students and the Scholarships for Research Students from the Ministry of Aliyah and Integration of Israel. D.U. is a Ph.D. student at the Faculty of Agriculture, Food, and Environment under the supervision of D.H. and C.G.

